# Time-resolved RNA-seq profiling of the infection of *Dictyostelium discoideum* by *Mycobacterium marinum* reveals an integrated host response to damage and stress

**DOI:** 10.1101/590810

**Authors:** N. Hanna, F. Burdet, A. Melotti, C. Bosmani, S. Kicka, H. Hilbi, P. Cosson, M. Pagni, T. Soldati

## Abstract

Tuberculosis remains the most pervasive infectious disease and the recent emergence of multiple drug-resistant strains emphasizes the need for more efficient drug treatments. The experimentally versatile *Dictyostelium discoideum – Mycobacterium marinum* infection model provides a powerful system to study mycobacteria pathogenicity and host response. In this study, a time-resolved transcriptomic analysis of the amoeba *D. discoideum* was performed to decipher the different host pathways impacted during infection. We investigated how *D. discoideum* fine-tunes its gene expression in response to *M. marinum* infection by assessing the transcriptomic profile covering the critical stages of entry, establishment of a permissive niche, proliferation and dissemination (1, 3, 6, 12, 24 and 48 hours post infection). Differential gene expression provided a fingerprint of the transcriptome of the host cell in the presence of mycobacteria, and helped identify specific markers and molecular signatures of infection. Enrichment pathway analysis showed that most of the Biological Processes (BP) of upregulated genes at early time point of infection hinted towards damage response and cellular defence, especially in specific pathways involved in membrane repair (ESCRT) and bacteria elimination (autophagy). Whereas at late time points of infection, BP related to starvation were upregulated. Some other signatures were more unexpected, such as cell cycle (downregulation of cytosolic large & small ribosomal subunits) and upregulation of metabolic adaptations (lipids transport).

## INTRODUCTION

Tuberculosis (TB) is an old but re-emerging global health threat caused by mycobacteria belonging to the *Mycobacterium tuberculosis* (Mtb) complex (WHO, 2018). About 95% of the affected individuals have a latent infection, in which Mtb is in a low metabolic, persistent and slowly or not replicating state. Nevertheless, ~10% of latent infections eventually progresses to active disease, which, if left untreated, kills more than half of the infected patients (Falzon et al., 2017). *Mycobacterium marinum* is a nontuberculous mycobacterial pathogen of fresh and salt water ectotherms. As with other nontuberculous mycobacteria (NTMs), *M. marinum* is an opportunistic human pathogen from environmental origin (Tobin and Ramakrishnan, 2008). Although *M. marinum* differs from *M. tuberculosis*, the two pathogens share virulence pathways that promote survival within the host. As such, *M. marinum* is a well-established and accepted model for several aspects of *M. tuberculosis* pathogenesis (Cardenal-Munoz et al., 2017b; Hagedorn and Soldati, 2007).

On the host side, *Dictyostelium discoideum* has become an important model organism to study the cell biology of professional phagocytes due to many shared molecular features with animal macrophages (Dunn et al., 2018). The broad range of existing genetic and biochemical tools, together with its suitability for cell culture and live microscopy, make *D. discoideum* ideal to investigate cellular mechanisms of cell motility, differentiation, and morphogenesis (Loomis, 2015; Nichols et al., 2015). Furthermore, *D. discoideum* possesses evolutionary conserved pathways involved in pathogen recognition and defences that are highly similar to those in macrophages (Bozzaro et al., 2008). In particular, thanks to the extreme conservation of phagosomal components and function with human phagocytic cells (Boulais et al., 2010), *D. discoideum* is used as a recipient cell to investigate interactions with pathogenic bacteria (reviewed in Dunn et al., 2017) (Hagedorn and Soldati, 2007; Solomon et al., 2003; Steinert and Heuner, 2005).

*M. marinum*, like Mtb induces phagosome maturation arrest, restricting its acidification and preventing evolution to a bactericidal milieu. Then, the phagosome becomes an active interface between the host cell and the mycobacteria, by interacting with host machineries such as autophagy (Cardenal-Munoz et al., 2017a), and compartments of endosomal origin and others (López-Jiménez et al., 2018). It finally loses its integrity, giving *M. marinum* access to the host cytosol (Cardenal-Munoz et al., 2017b). As a response, host cells developed strategies to fight against various pathogens. Protozoa use their intrinsic defences to recognize, contain or kill pathogens (Cosson and Soldati, 2008; Dunn et al., 2017). In addition, modulation of host cell processes such as autophagy (Songane et al., 2012), metabolism (Gouzy et al., 2014), ion homeostasis (Barisch et al., 2018; Botella et al., 2011; Bozzaro et al., 2013; Lefrancois et al., 2019; Soldati and Neyrolles, 2012) and programmed cell death (Briken et al., 2013) have been shown to participate in the clearance of bacteria.

In recent years, transcriptomic studies on mycobacteria during infection, *in vitro* and in whole organisms, resulted in major advances in our understanding of virulence strategies, and of how mycobacteria sense, respond and adapt to the intracellular and intravacuolar milieu (Keren et al., 2011; Liang et al., 2017; Rohde et al., 2012). Recent studies showed that Mtb has a unique transcriptional signature as part of its adaptive response to a hostile environment, involving metabolic reprogramming, respiration, oxidative stress, dormancy response, and virulence (Lin et al., 2016). Using the same genome-wide approach, the active transcriptome of neutrophils in zebrafish at the onset of *M. marinum* infection was studied, showing that neutrophils fight mycobacterial infection by an inflammasome-dependent mechanism during the initiation of infection (Kenyon et al., 2017). Many studies assessing the transcriptional response of the host or mycobacterium have been performed already, but few of them focused on the cross-talk between both organisms, and even less frequently covered the infection time-course in a high resolution manner (Betts et al., 2002; Lin et al., 2016; Rienksma et al., 2015).

Determining the transcriptome of intracellular pathogens during the infection time course remains difficult especially for mycobacteria due to the low bacteria-to-host RNA ratio. The load of intracellular bacteria varies between different pathogens, reaching few thousands for *Salmonella* and drops to less than 10 for Mtb (Rienksma et al., 2015; Westermann et al., 2016). In order to circumvent this problem, we used two separate enrichment strategies, one to determine the host transcriptome, and one to deplete host components to obtain the *M. marinum* transcriptome.

## RESULTS AND DISCUSSION

We aimed to recapitulate all the major events during the *M. marinum* infection time course so as to reveal how the host *D. discoideum* fine-tunes its gene expression in response to infection. Our time-resolved transcriptomic analysis of *D. discoideum* during infection, covered the critical stages of entry (1 hpi), establishment of a permissive niche (3, 6 hpi) vacuolar proliferation (12, 24 hpi), escape to the cytosol, further proliferation and dissemination (36, 48 hpi). Due to the low number of intracellular *M. marinum*, we performed a “parallel RNA-seq” to capture the various transcriptional strategies adopted both by *D. discoideum* [using a FACS-based enrichment strategy (figure 1 - red arrow)] and by *M. marinum* [using a GITC-based depletion strategy (figure 1 - green arrow)].

**Figure 1.**
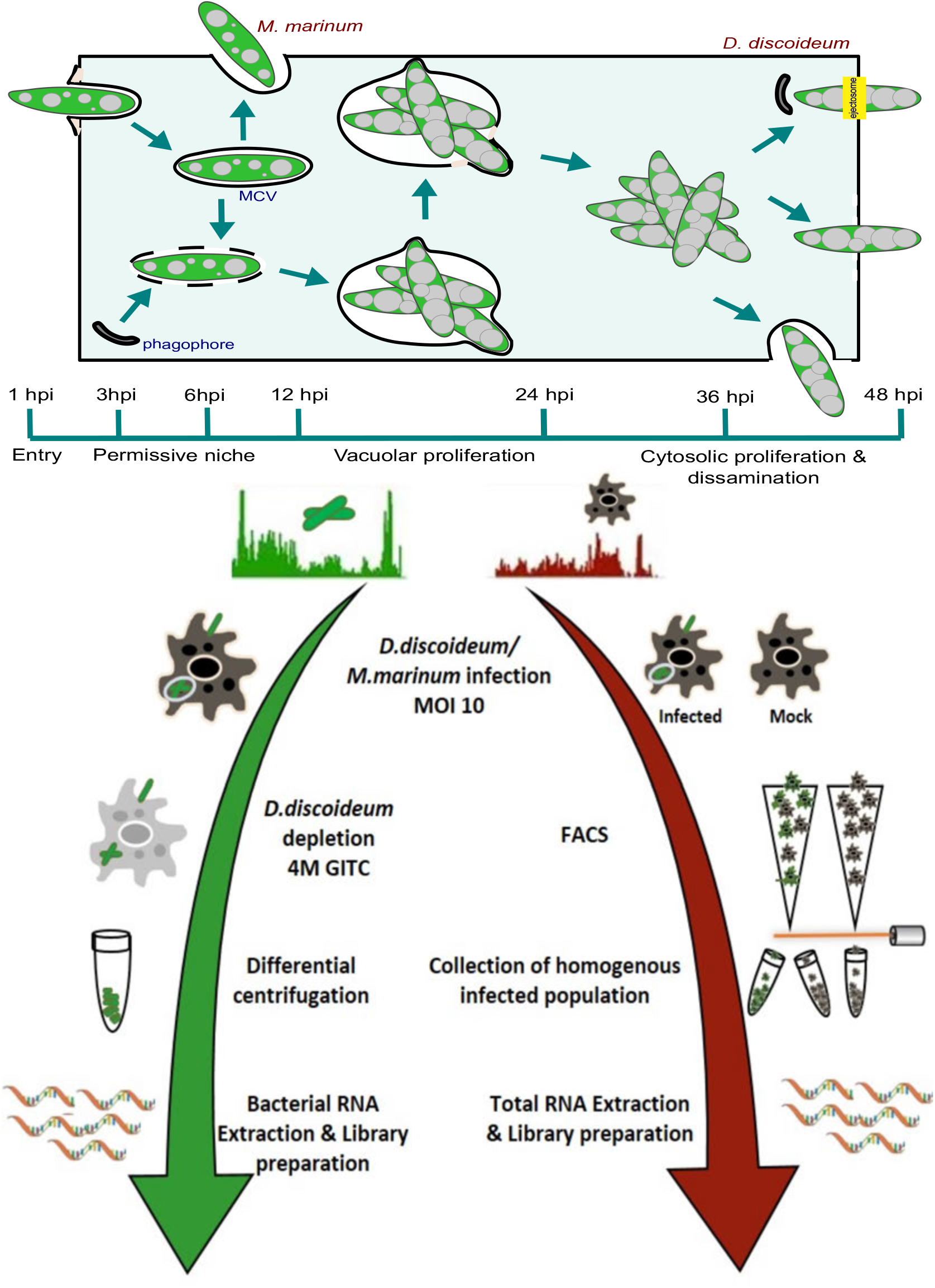
Workflow of parallel RNA-seq. *M marinum* stably expressing GFP were used to infect *D. discoideum* AX2 (ka) and samples were taken over a time-course covering the three aspects of the infection (early infection, establishment of permissive niche, and proliferation/dissemination). The collected samples contained a heterogeneous population of both infected (GFP-positive) and uninfected bystander (GFP-negative) cells. To obtain a homogeneous population of *M. marinum*-infected *D. discoideum*, the samples were subject to FACS. RNA-seq of mock-infected host cells served as a reference control (right Arrow). To obtain a high resolution *M marinum* intracellular transcriptome, an enrichment step was applied by depleting host material prior to analysis by guanidinium-mediated differential lysis of host cells (4 M GITC) (left Arrow).

### An optimized FACS protocol to isolate infected cells

We infected monolayers of adherent *D. discoideum* by spinoculating fluorescent *M. marinum* onto them, which ensured efficient and synchronous phagocytosis of the bacteria. A typical infection starts with 60% of infected cells at 1 hpi and drops to 10 % at 48 hpi due to infection dynamics and cell division. Therefore, cell sorting was needed to separate infected from non-infected cell subpopulations, and thus obtain the distinct and specific transcriptomes. The optimal cell sorting enabled separating host cells into three different subpopulations, infected cells (GFP+) and non-infected bystanders (GFP-) from the infection, as well as mock-treated cells (mock). We sorted on average 500,000 cells for each subpopulation and time point. The purity of the sorted populations was very consistent based on FACS reanalysis, indicating that almost 99% of sorted cells belonged to the correct subpopulation.

### lnDA-C primers efficiently removed rRNA of *D. discoideum*

cDNA libraries were prepared for RNA-seq using the Nugen protocol and rRNA was depleted using customized Dependent Adapter Cleavage (InDA-C), for effective removal of specific transcripts from RNA-Seq libraries without impacting non-targeted transcripts. We constructed strand-specific libraries using the Ovation kit (NuGen) and sequenced them using 50 bp single read Illumina chemistry. The 167 RNA-seq libraries used for the present study were sequenced across 21 lanes of an Illumina^®^ HiSeq 4000 sequencing apparatus. The cut-off for the acceptable number of reads per sample was 10 million reads after filtering for both adapter sequence and poor quality reads. Reciprocal mapping verified that no read from *D. discoideum* mapped to the genome of *M. marinum.* It has been shown that ribosomal RNA in *D. discoideum* constitutes more than 96% of the total cellular RNA, the data showed that our depletion strategy was successful by reducing the percentage of rRNA down to between 10% and 40 % of the total number of reads, consequently improving the coverage for the protein-coding mRNAs (figure S1). The amoeba *D. discoideum*, has an AT-rich (78%) genome of 34 Mb and about 12,257 protein-coding genes (Eichinger et al., 2005). Our transcriptome detected transcripts for 11,159 genes, providing comprehensive coverage under the conditions used in this study. For the downstream analysis, we kept 10,369 genes after filtering lowly expressed genes.

### Principal Component Analysis reveals clustering by the time post-infection

In order to explore the dynamics of gene expression throughout the infection time course, we used Principal Component Analysis (PCA), a widely used tool to investigate transcriptomic data sets. To improve the relevance and consistence of the clustering, the data were first corrected for date-batch effect, and then we excluded the genes with nonsignificant variation (cut-off p>0.05) before plotting all 3 subpopulations (GFP+, GFP-, and mock). The transcriptome of each sample was plotted in a twodimensional plane with axes corresponding to the first two principal components (PCs). PC1 and PC2 accounted for nearly half (47%) of the variation in the entire data set, whereas PC3 and PC4 explained on average another 14% of the variations (figure 2A, S2). Interestingly, the GFP+ samples clustered together but away from the GFP- and mock-treated, pointing towards profound differences induced by infection. Comparatively, GFP- and mock appeared grouped and thus more similar to each other on the PCA plot (figure 2B, 2C, 2D). To increase clarity and focus on the individual time points, new PCAs were calculated separately for each time point (figure S3) for the three different subpopulations (mock-treated, GFP+, and GFP-). The first two principal components explained between 34 and 68% of the total variance. A significant degree of similarity between biological replicates was evident in the PCA plots. Interestingly, mock-treated cells clustered tightly with uninfected cells at various time points. In contrast, the infected cells clustered together and distinctly away from other samples, suggesting an apparently unique *M.* marinum-specific *D. discoideum* response to infection. The global transcriptional patterns for the early time points were partitioned away from those observed at later time points (figure 2B). This was consistent with the pronounced and distinct host response to infection, which also displayed a clear separation between responses at early and later times post-infection (figure 3A, B).

**Figure 2.**
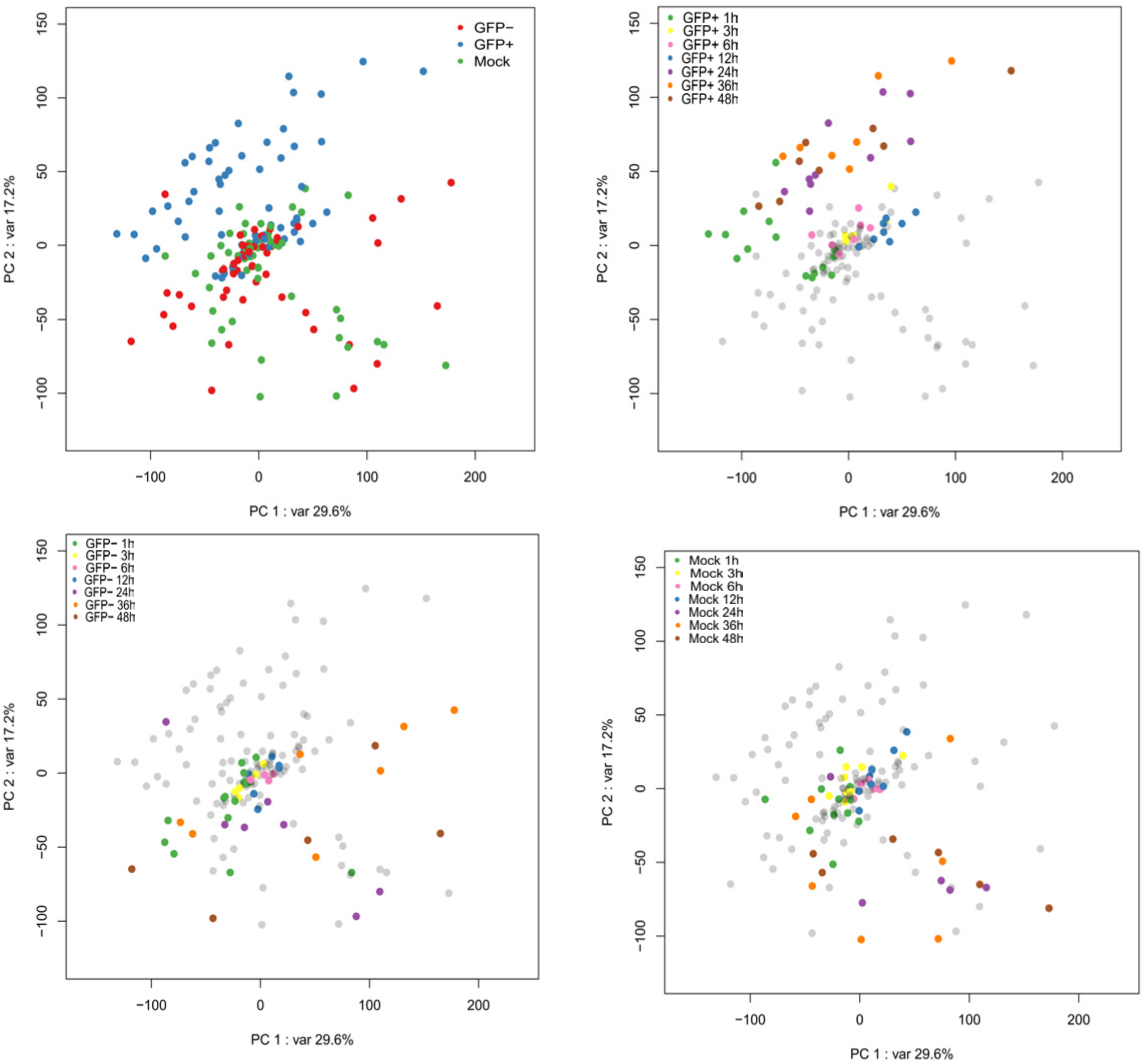
Principal component analysis. Each dot represents the gene expression profile of one sample representing a single subpopulation (A) or a single time point in a given condition (B, C, and D). The colours in A indicate the various experimental conditions: Mock-treated (Green), GFP+/infected (Blue), and GFP-/bystanders (Red). The axis labels indicate the percentages of explained variance corresponding to the represented principal component. All analyses were performed after filtering out non-significant expressed genes and including experimental batch as a covariate in the statistical model

**Figure 3.**
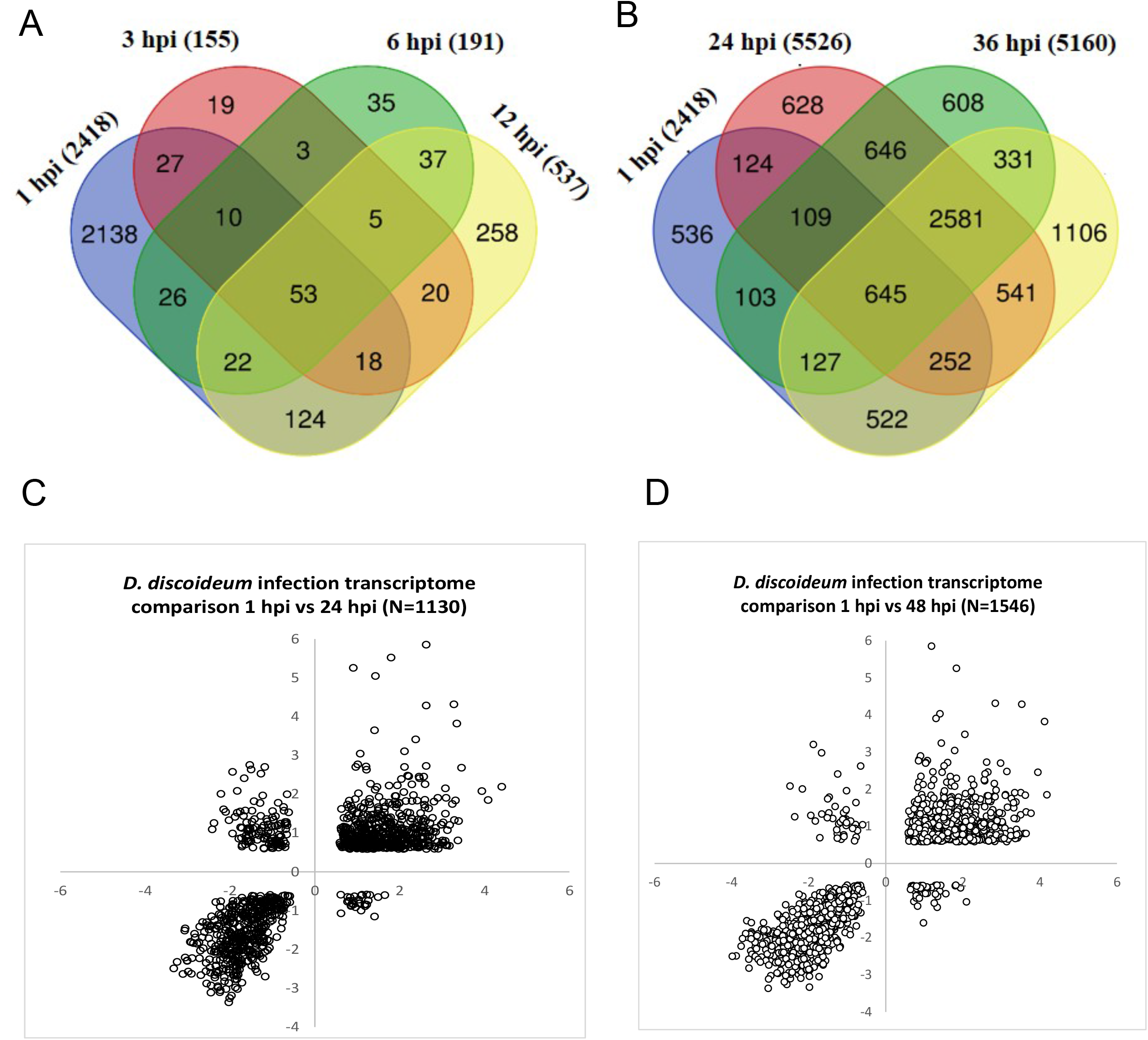
Temporal expression dynamics and Overview of whole-transcriptome data. (A) and (B) Venn diagrams showing the number of differentially expressed genes per comparison across time points and the overlap between each set of genes. Data for transcripts that were commonly differentially regulated (log_2_-fold change of ≥0.585, p-value ≤0.05) are shown. The total numbers of genes per time point are 2418 at 1 h, 155 at 3 h, 191 at 6 h, 537 at 12 h, 5526 at 24h, 5150 at 36h, and 6105 at 48 h. (C) and (D) correlation analysis of DE gene patterns at 1, 24, and 48 h. The gene clustering clearly shows a high correlation between the DE genes at these given time points (right upper and left lower quadrant).

### Analysis of differential gene expression shows an evolution of *D. discoideum* transcriptional response during the infection time course

The sequence reads that mapped to unique locations in the *D. discoideum* reference genome were used to generate lists of differentially-expressed (DE) genes between the *M.* marinum-infected and mock-treated control groups at all time points. Genes were considered to be differentially regulated if they showed, under at least one condition, an average transcript value with a log_2_ ratio greater than 0.585 or less than −0.585 with a p-value ≤ 0.05. DE genes in infected cells compared to their expression in mock-treated cells were identified by calculating reads per kilobase per million (RPKM). The analysis revealed a pronounced transcriptome change at the earliest infection time point (1 hpi, N= 2418), indicating a rapid remodelling of the transcriptome already at 1 h, in response to *M. marinum* pathogenicity and the resulting membrane damage (López-Jiménez et al., 2018). The number of DE genes drastically dropped during the time corresponding to the formation of a permissive niche (3 hpi, N= 155 & 6 hpi, N=191), giving rise to a new molecular signature, followed by another peak of transcriptomic changes (12 hpi, N =537). At 1 and 12 hpi the number of DE genes were almost evenly distributed between upregulated (up) (294 and 1029 genes, respectively) and downregulated (down) (243 and 1389 genes, respectively) (Table S1). It is important to note that the number of DE genes observed between infected and mock-treated samples at 3 and 6 hpi was markedly higher for upregulated genes (123 and 129) compared to the downregulated genes (32 and 62). Interestingly, most of the DE genes at 3 and 6 hpi were common with the other 2 early time points leaving only few time point-specific genes (19 and 35 genes at 3 and 6 hpi, respectively) (figure 3A). Surprisingly, during the progression of infection, the number of DE genes at later time points increased enormously reaching more than 60% of the total monitored transcriptome (48 hpi, N= 6105) Notably, the difference between the number of up and down DE genes was not as marked at the 24 hpi (2846 up and 2680 down) and 48 hpi (3137 up and 2968 down) compared to 36 hpi (3354 up and 1796 down), when the number of downregulated genes was lower by almost 25%. This data suggests that the stages of infection corresponding to proliferation of *M. marinum* and its escape to the cytosol has a greater effect on the host cell compared to the stage of establishment of a permissive niche.

To investigate the regulatory patterns of DE genes, we plotted Venn diagrams of early and late time points separately, keeping the 1 hpi as an anchor point for both comparisons. This analysis revealed a core of only 53 DE genes that were identified throughout the early infection course (figure 3A). This number was substantially higher at the late time point reaching 645 DE genes that are either up or downregulated (figure 3B). These 2 values constitute 2.2% and 26.7% of the DE genes identified at 1 hpi, respectively. This comparison clearly shows that this time-resolved transcriptome is very dynamic and involves many key players that varies throughout the different stages of infection. Taken all together, we identified 22 genes differentially expressed during the whole infection time-course which can be potentially considered as global infection markers. Among them, VacC, one of the three flotillin homologues in *D. discoideum* that is involved in the recycling of plasma membrane proteins and susceptibility to infection (Bosmani et al., 2019), the superoxide dismutase, SodB and a protein with a Ubiquitin-binding CUE domain (DDB_G0270388). We next examined the correlation between the log_2_ expression values generated by comparing the transcriptome *of D. discoideum* at 1, 24 and 48 hpi for all DE genes that passed the filtering criteria. We noticed a regular trend of the DE at these time points with very few genes showing opposite expression pattern (figure 3C, 3D). The substantial overlap in DE genes as well as their conserved expression trend, suggests that the *D. discoideum* transcriptional response to *M. marinum* has a consistent pattern of regulation. An in depth inspection of DE genes using volcano plots showed a clear trend as a function of infection progression, in which the amplitude of genes that are upregulated is higher than the ones that are downregulated (figure 4). Markedly, the top upregulated gene had a 64-fold increase (*iliL*) compared to 27-fold decrease for the most downregulated genes (DDB_G0272094). Interestingly, while the number of up and down DE genes is quite comparable, the volcano plots clearly visualise the higher number of highly-significantly upregulated genes compared to the downregulated ones (figure 4A, E, and G). It is noteworthy that the downregulated DE genes are separated into two groups on the volcano plots especially at the late time points of infection (figure 4 F, G).

**Figure 4.**
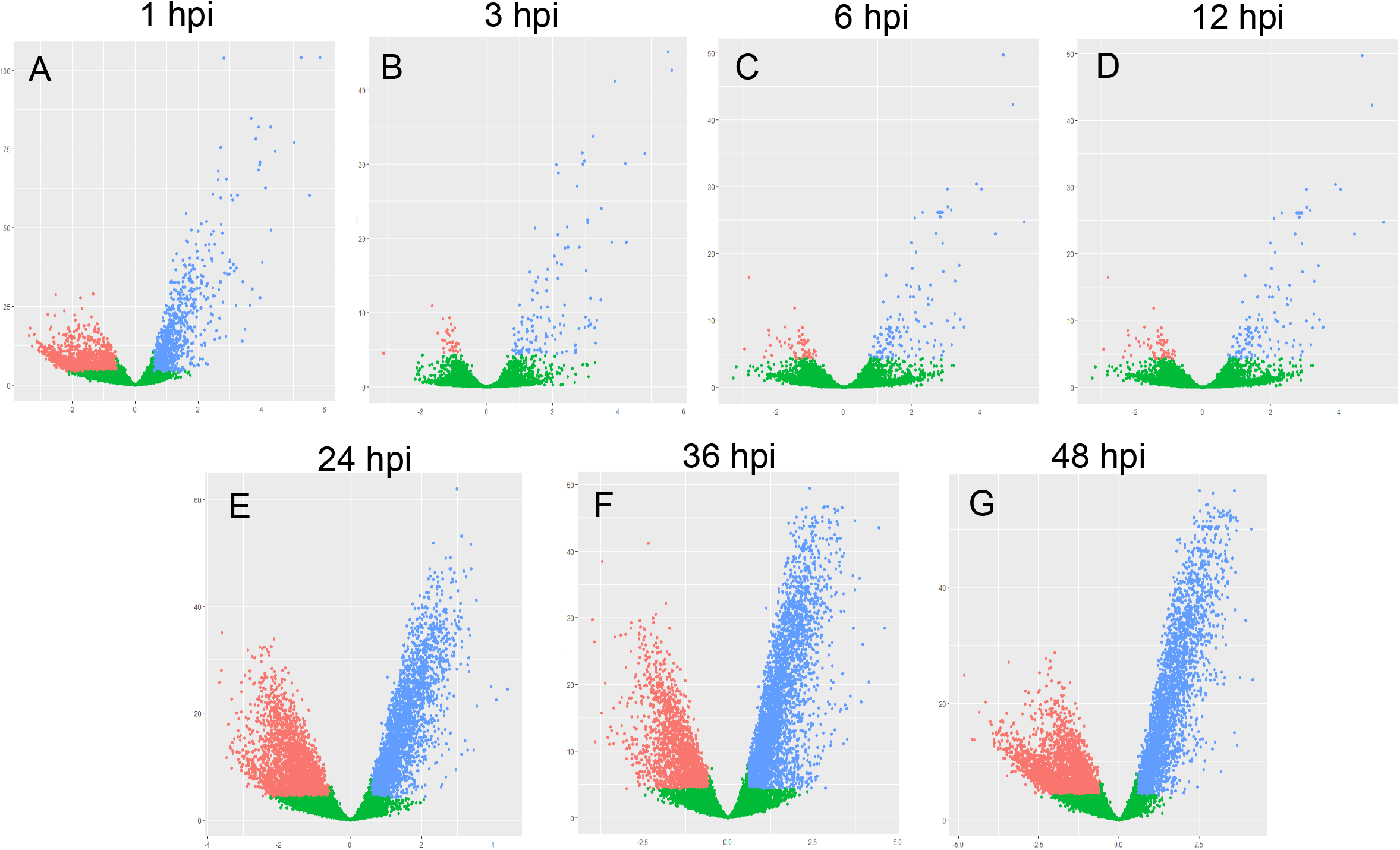
Summary of the RNA-seq results. Volcano plot representation of differential expression analysis of genes in the infected versus mock-treated comparison for the various time points (A=1, B=3, C=6, D=12, E=24, F=36, G=48 hpi). Blue and red dots mark the genes with significantly increased or decreased expression respectively in infected compared to Mock samples (p-Value ≤0.05). The x-axis shows log_2_ fold-changes in expression and the y-axis the p-value (log10) of a gene being differentially expressed.

### Pathway-enrichment analysis depicted a multifaceted host response to infection

An important aim of this study was to define molecular signatures of *D. discoideum*’s response to the modulation of the intracellular environment by *M. marinum.* To reveal the general trends of these responses, we performed a TopGO enrichment analysis of Biological Processes (BP). To more finely appreciate the pathways that are stimulated or repressed. DE genes were separated in up and down categories for the analysis (see table S2). BP appeared to be relatively consistent between the different postinfection time points and are, for the most part, associated with host defence mechanisms (host defence response, proteasome assembly, phagosome maturation, and phagocytosis), which were upregulated, while some cellular signalling (small GTPase mediated signal transduction and G-coupled protein receptors) was downregulated (figure 5). The transcriptome signatures at 1 hpi showed that transcription and RNA modification were downregulated on a large scale, clearly indicating that *D. discoideum* senses *M. marinum* infection as a stress or a starvation-like condition (figure 5A and B). The latter two BP groups were again detected during the stage of establishment of a permissive niche (3 and 6 hpi), in addition to cell division and DNA replication which were also downregulated (figure 5C, D, E, F). At 12 hpi, groups related to GTP biosynthesis as well as mitotic spindle were downregulated. At the late infection time points, mitochondrial transport, microtubule-based movement, and actin nucleation were downregulated (Figure 6A, C, E). Whereas, signaling-related BP groups were upregulated including phosphatidylinositol phosphorylation, Rho and other GTPases and regulators (figure 6 B, D, F), which indicates a global transcriptional reprogramming and possibly explains the drastic increase in the number of DE genes at these time points. Notably, most of the BPs of upregulated genes at early time point of infection hinted towards damage response and cellular defence, compared to BP related to transcriptional reprogramming and starvation that were upregulated at late time points.

**Figure 5 and 6.**
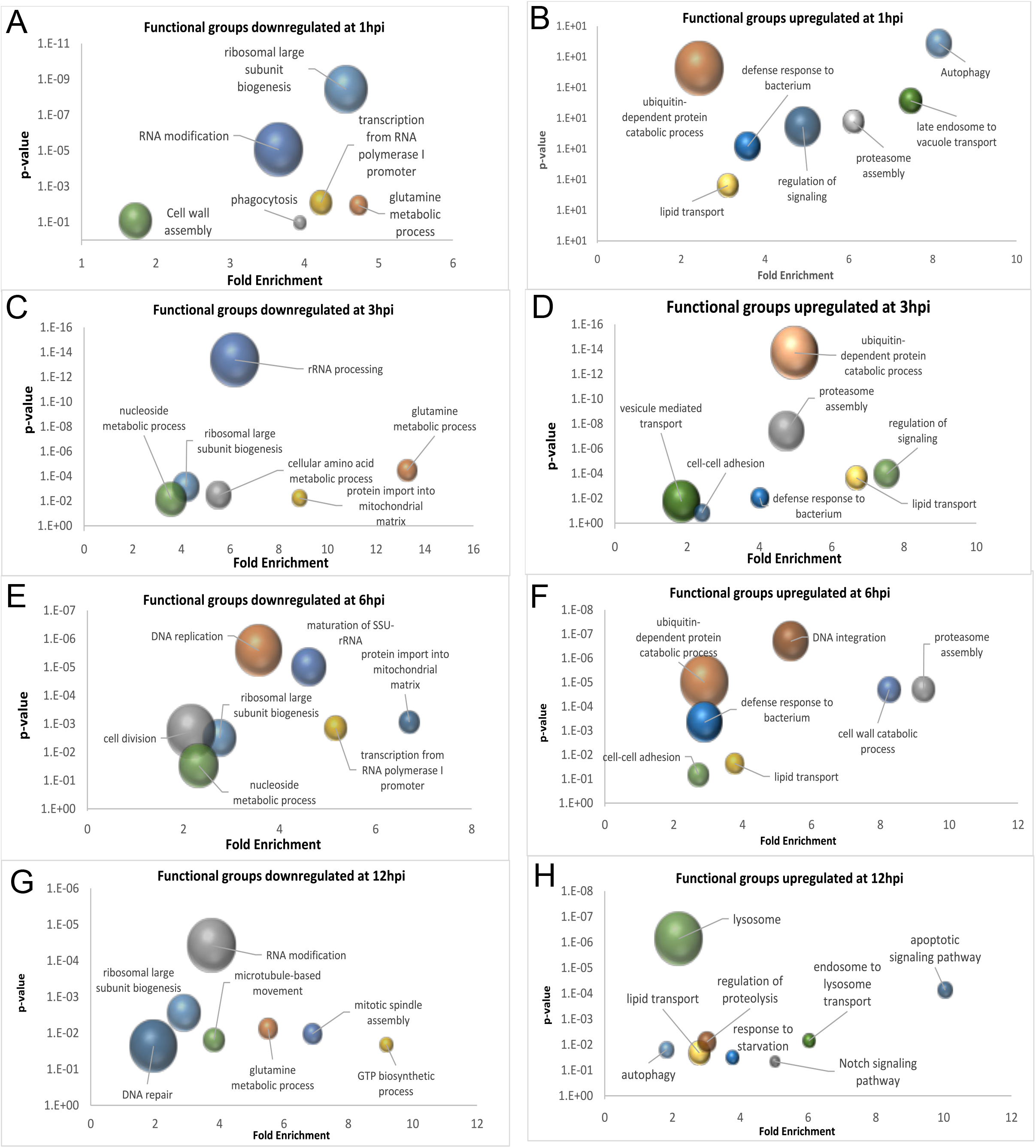
TopGO pathway analysis. TopGO analysis was carried out to identify pathways related to biological processes overrepresented in genes that constitute *D. discoideum* response to *M. marinum* infection relative to mock controls. Genes that were differentially expressed (DE) by more than 1.5-fold were used as input with up- and down-regulated genes considered separately. For each pathway, the fold enrichment score (x axis) was plotted against its p-value (y axis). The number of DE genes assigned to that pathway is presented by the size of the bubble.

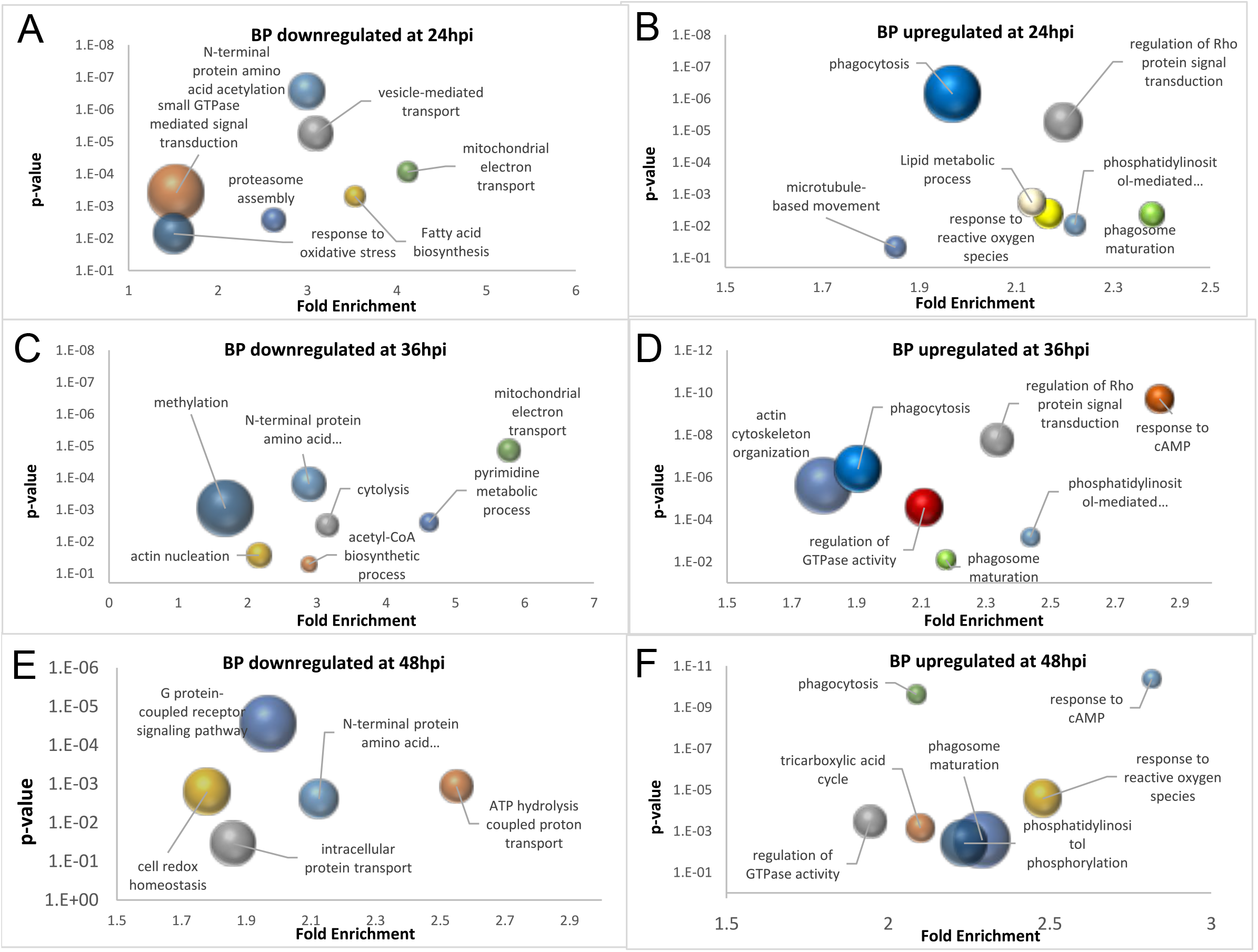

### *M. marinum* infection induces a damage and stress response in *D. discoideum*

In order to gain more specific insights about the key players during the infection, we performed a Gene Set Enrichment Analysis (GSEA) which is complementary to single-gene approaches, and provides a framework to examine changes operating at a higher level of biological organization. In this analysis, the ranking of genes based on their expression in a predefined set of genes that can be found at the molecular signatures database (MSigDB) (Liberzon et al., 2015). More precisely, during the early infection time points, pathways that were most enriched in the upregulated sets of genes included components of cell-autonomous defences, and especially mainly genes related to autophagy, generation of reactive oxygen species (ROS) and phagocytosis as well as ubiquitin ligases (figure 7 AD). In parallel, gene sets related to transcription were downregulated, mainly polymerase I and II subunits and small and large ribosomal subunits proteins. Additionally, the late time points showed a clear downregulation of lytic enzymes and ATP hydrolysis-coupled proton transport and an upregulation of genes belonging to the TOR pathway (regulation of the intracellular signal transduction). Lipid metabolism (ceramide synthases) and transport (ABC transporters) seem to play a central role during the infection time-course as genes belonging to these groups were upregulated at various time points, which is in perfect agreement with the documented interface between the host and the pathogen lipid metabolisms (figure 7 E-G) (Barisch et al., 2015; Barisch and Soldati, 2017).

**Figure 7:**
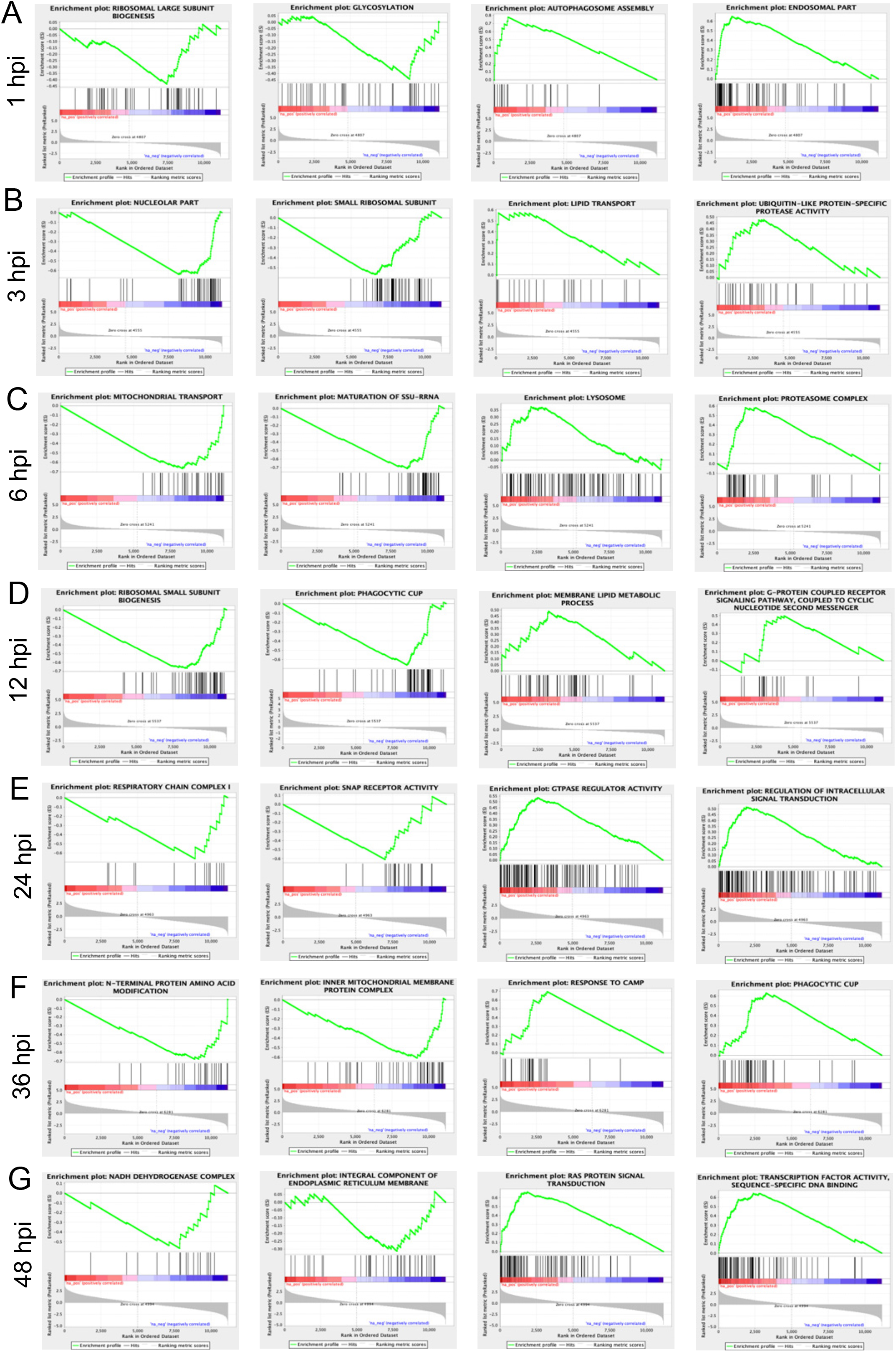
GSEA analysis of the differentially expressed genes during *D. discoideum* infection. Example GSEA enrichment plots for selected gene sets differentially expressed at the different time points. On the x axis are genes ranked according to their expression, from the up-regulated genes (positively regulated) on the left hand side to the down-regulated genes (negatively regulated) on far right. Black vertical lines show the positions of the individual genes in the gene set. The cumulative value of the enrichment score (y axis) is represented by the green line. A positive normalized enrichment score (NES) indicates enrichment in the up-regulated group of genes, while a negative NES indicates prevalence of the genes in the down-regulated group. The title for each of the graphs indicates the gene set used in the comparison.

During the infection course, we identified an early response consisting in the upregulation of transcription of a large collection of phagocytosis and lysosome related genes (phg1A, AlyA, B, C, D) (Perrin et al., 2015). In addition, consistent with the report that ESCRT plays an important role in membrane repair during *M. marinum* infection, many of the ESCRT components were induced (López-Jiménez et al., 2018). Genes coding for autophagy-related proteins such as Atg1, the two paralogs Atg8a and Atg8b, and p62/SQSTSM1 among others were also upregulated through the whole infection time course (see Table S1)(Cardenal-Munoz et al., 2017a). In parallel, other host defence genes were also upregulated, notably *noxB*, involved in ROS generation, was increased more than 8-fold at the late infection time points (Zhang et al., 2013). A prominent defence mechanism identified in *D. discoideum*, used to kill bacteria (Barisch et al, 2018) and restrict an infection by *M. marinum* (Lefrancois et al., 2019), is metal intoxication. Consistent with this mechanism, the zinc transporters ZntA and ZntB were upregulated at the three stages of infection. More unexpectedly, but confirming the GO term enrichment analysis, we also obtained signatures of cell cycle regulation, with a consistent downregulation of the cytosolic large and small ribosomal subunits at early time points of infection, including genes such as rrs1, a regulator of ribosome biogenesis. This molecular signature potentially indirectly indicates a post-translational downregulation of the TOR signalling pathway. Also, at the late infection stage, many vacuolar ATPase subunits were downregulated (Vat C, D, E, F, G, H), which is consistent with previous knowledge of the interference of *M. marinum* with phagosome maturation (Koliwer-Brandl et al., 2019; Lerena and Colombo, 2011). The late time point also brought to light an unexpected molecular signature involving upregulation of genes encoding many components of the TOR pathway, including Tor, Raptor, RipA, and several kinases (PkgA, B, and IkA, B, C). A number of studies have shown the cross talk between TOR and autophagy and have identified the main key players of this interaction, but so far very little is known about this cross-talk during infection (Bakula et al., 2017; Dooley et al., 2014).

## MATERIALS AND METHODS

### *D. discoideum* and mycobacteria strains, culture, and plasmids

*D. discoideum* Ax2(Ka) was cultured axenically at 22°C in Hl5c medium (Formedium) supplemented with 100 U mL-1 of penicillin and 100 μg mL-1 of streptomycin (Invitrogen). *M. marinum* (M strain) wt was cultured in 7H9 (Difco) supplemented with 0.2% glycerol (v/v) (Biosciences), 0.05% Tween-20 (v/v) (Sigma) and 10% OADC (v/v) (Middlebrock). GFP-expressing bacteria were obtained by transformation with msp12::GFP vector, and cultured in the presence of 50 μg/mL-1 kanamycin (AppliChem).

### Infection assay

Infections were performed as previously described (Kicka et al., 2018). In brief, GFP-expressing *M. marinum* grown overnight were centrifuged and re-suspended in Hl5c, and then spinoculated on adherent *D. discoideum* cells. Infections were performed at a multiplicity of infection (MOI) of 10. After washing off uningested bacteria, the infected cells were re-suspended to a density of 1 x 10^6^ cells/mL in filtered HL5-C. 5 μg/mL of streptomycin and 5 U/mL of penicillin were added to prevent extracellular bacteria growth. The infected cells were then incubated at 25°C at 130 rpm, and samples were taken for analysis at 1, 3, 6, 12, 24, 36, and 48 hpi. To control for changes in gene expression induced by culture conditions and treatments, a non-infected yet mock-treated control was included as a reference.

### Flow cytometry and fluorescence-activated cell sorting (FACS)

In order to obtain a specific and homogenous host response, and to enrich for bacterial RNA, infected and uninfected bystander cells were separated using flow cytometry-based live sorting. *D. discoideum* cells infected with *M. marinum-GFP* or mock-treated were collected, centrifuged (5 min at 1500 rpm) and re-suspended in HL5c. Cells were sorted using an Astrios device (Beckman), at 4°C by cooling both the input tube holder and the collection tube rack. Selection was achieved by gating on cell diameter (forward-scatter) and granularity (side-scatter). The infected (GFP-positive, GFP+) and non-infected (GFP-negative, GFP-) sub-fractions were defined based on GFP signal intensity (FITC channel) versus auto-fluorescence (PE channel). The gates for GFP+ and GFP-fractions were conservative in order to prevent cross-contamination. The same sorting settings and parameters were used for mock-treated cells. Typically, 5×10^5^ cells of each fraction were collected, centrifuged and re-suspended in TRIreagent (product nb. T9424, Sigma-Alrich) for RNA isolation (Westermann et al., 2017).

### Biochemical depletion of host components

Infections were performed as described above. Enrichment of mycobacteria from infected *D. discoideum* cells was carried out by differential lysis of host and mycobacterial cells by guanidine thiocyanate (GITC) (Chomczynski and Sacchi, 1987). The infected cells were re-suspended and lysed using 4 M cold GITC, and then transferred to a 1.5-ml Eppendorf tube. The sample was centrifuged for 1 min at 4°C/13,000 rpm and the supernatant was transferred to a fresh tube for RNA isolation.

### RNA Isolation and QC analysis

RNA was extracted from cells using the directzol RNA extraction kit (Zymo research) following the manufacturer’s instructions for total RNA isolation. Such a lysis buffer fails to break the thick envelope of *Mycobacterium*, so an additional step was added which consists of mechanically breaking the cells with beads (FastPrep Instrument). To ensure consistence, all the samples, including infected, uninfected and mock-treated cells were disrupted by bead beating (FastPrep Instrument; two cycles of 30 s at maximum speed with cooling on ice between cycles). To remove contaminating genomic DNA, samples were treated with 0.25 U of DNase I (Zymo) per 1μg of RNA for 15 min at 25 °C. RNA was quantified using Qubit 4.0 (Invitrogen) and its quality was checked on the Agilent 2100 Bioanalyzer (Agilent Technologies).

### Construction of cDNA libraries and RNA sequencing

Total RNAs, either from *M. marinum* depleted of host, or from *D. discoideum* alone, or as a mixture of D. *discoideum* and *M. marinum* RNA (in the case of infected cells), were subjected to cDNA synthesis and NGS library construction using the Ovation Universal System (NuGEN Technologies, San Carlos, California, USA). 100 ng of total DNAse I-treated RNA was used for first- and then second-strand cDNA synthesis following the manufacturer’s protocol. In order to obtain comparable library size, a double bead cut strategy was applied using the 10X genomics protocol. cDNA was recovered using magnetic beads with two ethanol washing steps, followed by enzymatic end repair of the fragments. Next, barcoded adapters were ligated to each sample, followed by an enzymatic strand selection step and magnetic bead recovery, as above. rRNAs were targeted for depletion by the addition of custom designed oligonucleotides specific for *D. discoideum* and *M. marinum* rRNAs (5S,18S, 28S and 5S, 16S, 23S, respectively). To amplify the libraries, 18 cycle of PCR were performed based on QC experiments carried out using RT-PCR.

The quality of the libraries was monitored by TapeStation (Agilent, High Sensitivity D1000 ScreenTape, # 5067–5584). Six-plexed samples were pooled in approximately equimolar amounts and run in 50bp single read flow cells (Illumina, # 15022187) and run on a Hiseq 4000 (Illumina).

### Bioinformatic analysis

RNA-seq libraries from infected and mock-treated cells taken at the indicated time points were analysed in pairwise comparisons. 50 nt single-end reads were mapped to the *D. discoideum* genome (downloaded from dictybase) (Fey et al., 2008) using tophat (version 2.0.13) and bowtie2 (version 2.2.4) softwares. The same procedure was applied to the bacteria-enriched samples and the reads were mapped to the *M. marinum* genome (M strain) (Stinear et al., 2008). As the RNA-seq data is stranded, parameter library-type was set to fr-second strand. Multi hits were not allowed, by using option --max-multi hits 1. The other parameters were set to default. The read counts per gene were generated using HTSeq software (version 0.6.1) and the GFF annotation downloaded from dictybase (February 2019). Options for htseq-count were -t exon --stranded = yes -m union. The counts were then imported in R (version 3.2.2). The genes were filtered for minimal expression, by removing genes with an average through all samples lower than 5 reads. Normalization factors to scale the libraries sizes were calculated using edgeR. The read counts were then log-transformed and variance stabilized using voom. The log-transformed counts were then batch-corrected for date effect using the R package limma and the removeBatchEffect function.

A differential expression analysis was then using the R package limma, including the date batch effect in the design. In total 14 comparisons were performed between the 2 conditions for the 7 time points. The genes having an adjusted p-value lower than 0.05 and an absolute fold change above 1.5 were considered differentially expressed. The union of these genes was then taken for the following analyses. The principal component analysis was generated using the R function prcomp, with centering and scaling of the data. The first 4 principal components were considered and plotted versus each other.

### TopGO (GO) Analysis

GO categories enriched in the *D. discoideum* DE gene lists were identified using the topGO package in R. For each comparison, upregulated and downregulated gene sets were input separately into topGO. A p-value cutoff of 0.05 was used. First, the weight01 algorithm was used to get the lowest level significant terms for each comparison. Then the union of these terms was used to run the classic Fisher algorithm, in order to be able to compare the results between all the comparisons.

### GSEA Analysis

The GSEA software from the Broad Institute (V3.0) was run in command line, using the rank lists from the limma differential analysis and the GO annotation from dictybase. We used the weighted algorithm, 1000 permutations and discarded the gene sets smaller than 15 genes and bigger than 500 genes.

## ACKNOWLEDGEMENTS

We gratefully acknowledge the staff of the FACS core facility and the iGE3 Genomics platform at the Faculty of Sciences and Faculty of Medicine of the University of Geneva for their precious help. This work was supported by an RTD grant from SystemsX.ch (awarded to TS, PC, HH, MP) and multiple grants from the Swiss National Science Foundation to TS. TS is also a member of iGE3.

---

**Figure S1.**
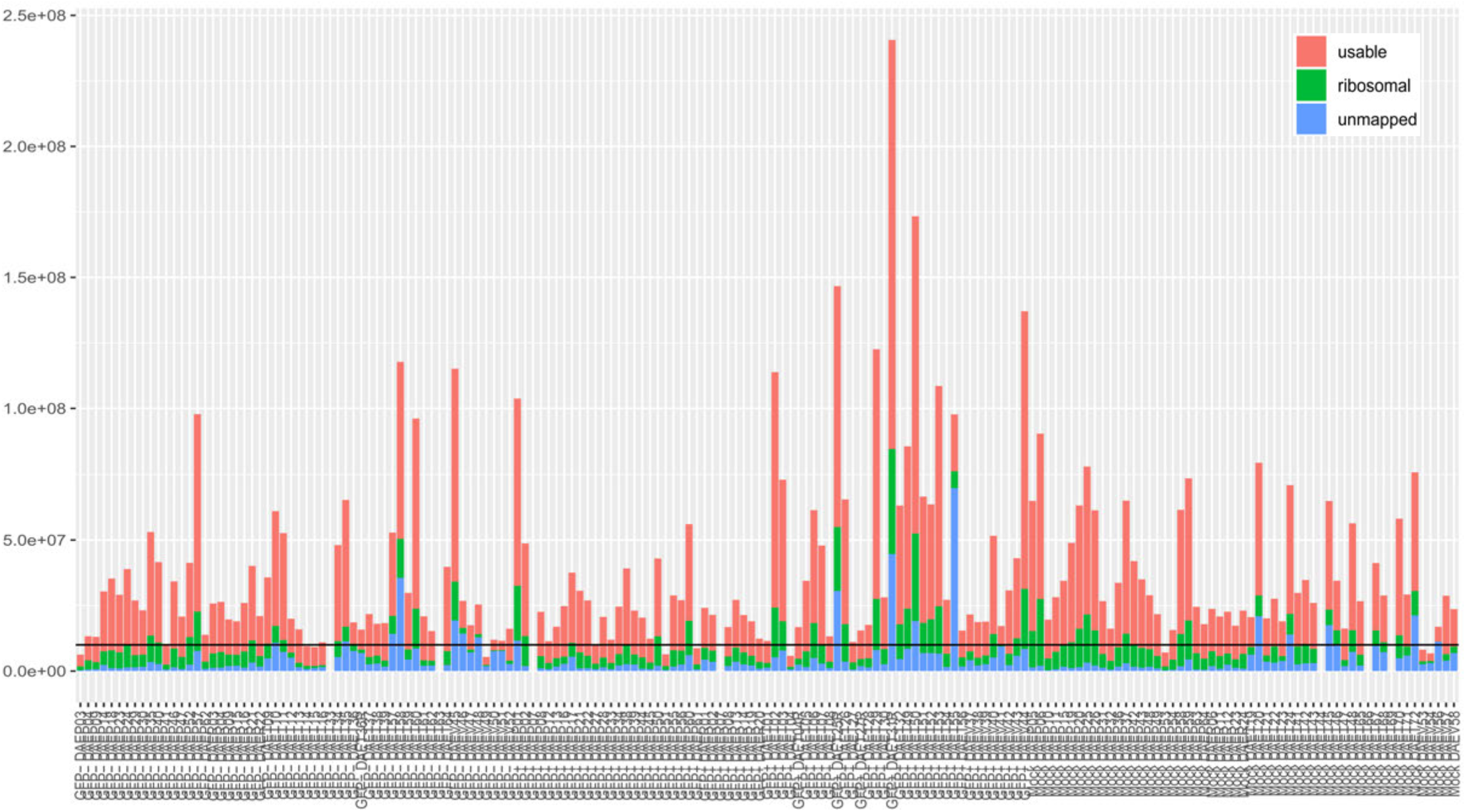
Efficient transcript depletion with AnyDeplete (lnDA-C). The plot shows the three types of obtained reads: usable (red), ribosomal (green), and unmapped (blue). All the samples included in the analysis passed the cut off value (10 M usable reads).

**Figure S2.**
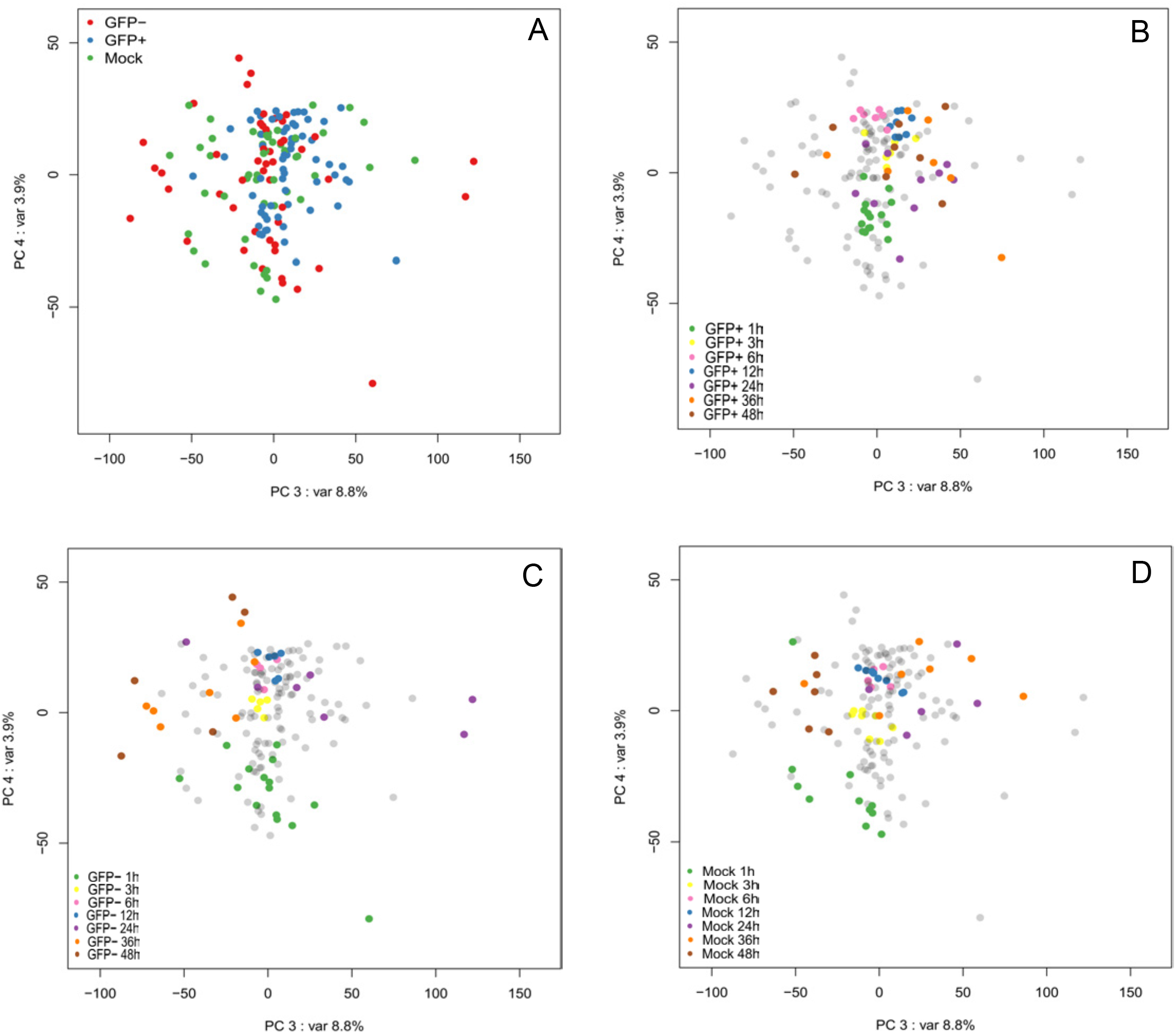
Principal component analysis. The axes correspond to the third (x axis) and the fourth (y axis) principal components. The samples descriptions are identical to figure 2.

**Figure S3.**
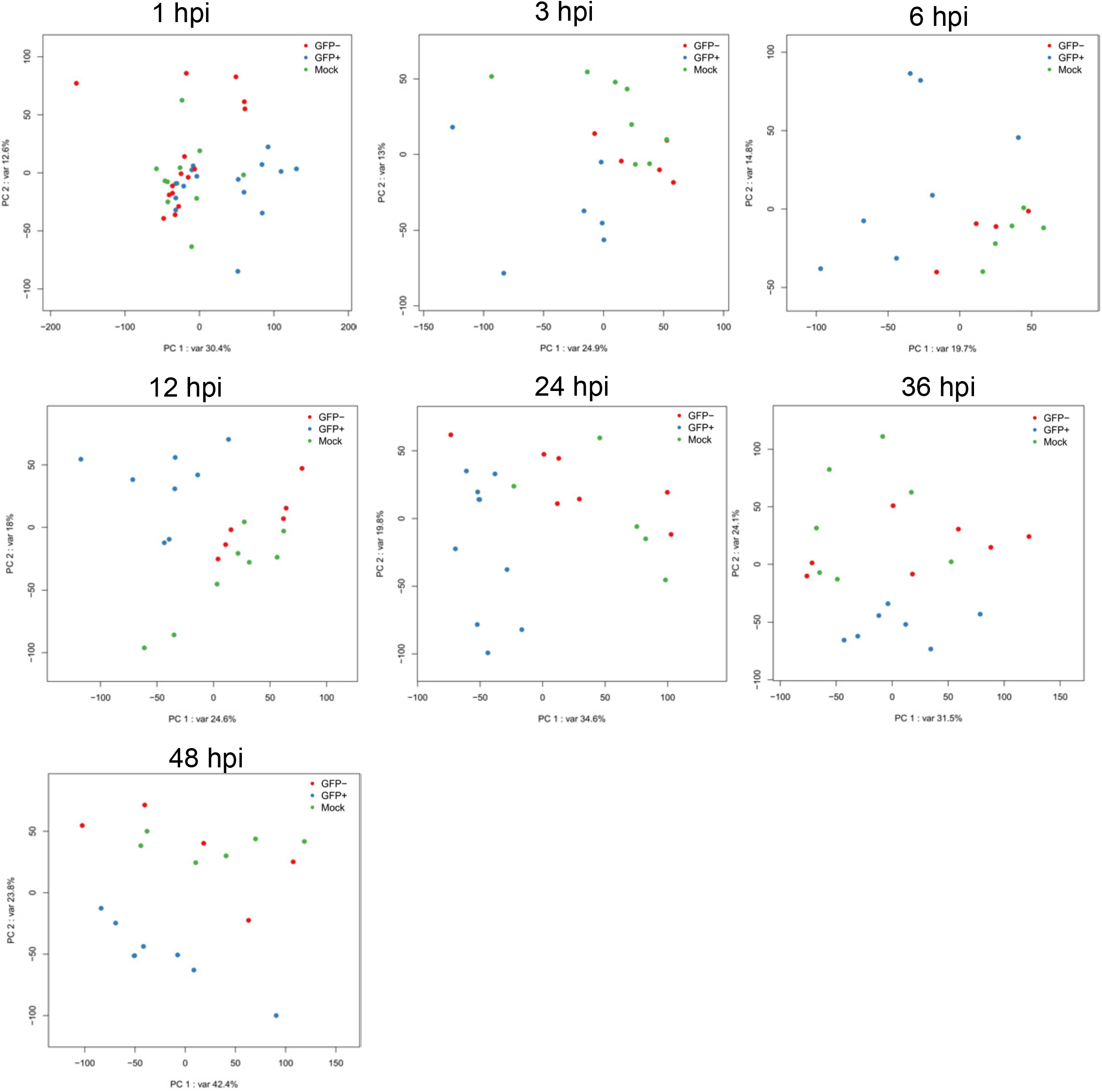
Principal component analysis. Each dot represents the gene expression profile of one sample representing a single subpopulation at a given time point. The sample description is identical to figure 2.

**Sup. Table 1.** List of differentially Expressed genes identified in the time resolved RNA-seq transcriptome.

**Sup. Table 3.** List of BP groups Identified by TopGo analysis

**Sup. Table 3.** List of enriched groups identified by GSEA.

